# Murine cytomegalovirus infection of melanoma lesions delays tumor growth by recruiting and re-polarizing monocytic phagocytes in the tumor

**DOI:** 10.1101/597948

**Authors:** Nicole A Wilski, Christina Del Casale, Timothy J Purwin, Andrew E Aplin, Christopher M Snyder

**Affiliations:** Department of Microbiology and Immunology, Thomas Jefferson University, Philadelphia, PA19107; Department of Cancer Biology, Thomas Jefferson University, Philadelphia, PA19107; Sidney Kimmel Cancer Center, Thomas Jefferson University, Philadelphia, PA19107

## Abstract

Cytomegalovirus (CMV) is a ubiquitous β-herpesvirus that infects many different cell types. CMV has been found in several solid tumors and it has been hypothesized that it may promote cellular transformation or exacerbate tumor growth. Paradoxically, in some experimental situations, CMV infection delays tumor growth. We previously showed that wild-type murine (M)CMV delayed the growth of poorly immunogenic B16 melanomas via an undefined mechanism. Here we show that MCMV delayed the growth of these immunologically “cold” tumors by recruiting and modulating tumor-associated macrophages. Depletion of monocytic phagocytes with clodronate completely prevented MCMV from delaying tumor growth. Mechanistically, our data suggest that MCMV recruits new macrophages to the tumor via the virus-encoded chemokine MCK2, and viruses lacking this chemokine were unable to delay tumor growth. Moreover, MCMV infection of macrophages drove them toward an M1-like state. Importantly, adaptive immune responses were also necessary for MCMV to delay tumor growth as the effect was substantially blunted in Rag-deficient animals. However, viral spread was not needed and a spread-defective MCMV strain was equally effective. In most mice, the anti-tumor effect of MCMV was transient. Although the recruited macrophages persisted, tumor regrowth correlated with a loss of viral activity in the tumor. However, an additional round of MCMV infection further delayed tumor growth, suggesting that tumor growth delay was dependent on active viral infection. Together, our results suggest that MCMV infection delayed the growth of an immunologically “cold” tumor by recruiting and modulating macrophages in order to promote anti-tumor immune responses.

**Importance:** Cytomegalovirus (CMV) is an exciting new platform for vaccines and cancer therapy. Although CMV may delay tumor growth in some settings, there is also evidence that CMV may promote cancer development and progression. Thus, defining the impact of CMV on tumors is critical. Using a mouse model of melanoma, we previously found that murine (M)CMV delayed tumor growth and activated tumor-specific immunity, although the mechanism was unclear. We now show that MCMV delayed tumor growth not by infecting and killing tumor cells, but rather by recruiting macrophages to the tumor. A viral chemokine was necessary to recruit macrophages and delay tumor growth. Furthermore, MCMV infection altered the functional state of the macrophages. Finally, we found that repeated MCMV injections sustained the anti-tumor effect suggesting that active viral infection was needed. Thus, MCMV altered tumor growth by actively recruiting and infecting macrophages in the tumor.

## Introduction

Cytomegalovirus (CMV) is a ubiquitous β-herpesvirus that establishes a systemic latent/persistent infection. While CMV can cause severe disease as a congenital infection or in immune compromised patients^1,2^, infection in healthy individuals is typically asymptomatic and is characterized by incredibly robust immune responses, including the accumulation of very large, sustained T cell populations^3–9^. Moreover, our previous data has suggested that even spread-defective variants of CMV may be able to sustain these T cell populations^10^. For these reasons, our group and others have been exploring the use of CMV as a vaccine platform for cancer and infectious diseases^11–22^.

Exploiting viruses and other pathogens in order to alter the tumor-microenvironment and enhance immune responses has garnered increased attention since the FDA approval of T-VEC in 2015^23^. T-VEC, the most successful oncolytic virus to date, is based on herpes simplex virus 1 and is currently being used as an intratumoral (IT) treatment for melanoma in the US, Europe, and Australia^24,25^. An alternative group of IT immune therapies in earlier stages of development are not designed to directly kill tumor cells, but rather to alter the microenvironment. These include Toxoplasma gondii, Salmonella, Semliki Forest virus, and inactivated modified Vaccinia virus Ankara, all of which have been tested pre-clinically in a B16 melanoma model and all of which function by altering the cytokine milieu, increasing leukocyte infiltration of tumors, and/or increasing antigen recognition or function of CD8+ T cells^26–32^.

Although early pre-clinical data suggest that CMV-based vaccines may have promise in models of prostate cancer, melanoma and squamous cell carcinoma^16–22^, it has also been proposed that latent CMV infection of tumors may contribute to tumor progression in patients^33–35^. Indeed, CMV has been found in several human cancers, with unknown consequences^36–40^. The best-studied association of cancer and CMV is with glioblastoma, in which anti-viral therapy may delay tumor growth^41–45^. Recent evidence suggested that murine (M)CMV may promote blood flow and angiogenesis in glioblastoma mouse models^46^. Additionally, previous work suggested that MCMV could promote rhabdomyosarcomas in mice lacking one copy of p53^47^. In contrast, several studies suggest that CMV may have the opposite effect and block tumor growth in other situations. Work from the Reddehase lab suggested that CMV infection may delay the growth of lymphoma in the liver^48–50^. Likewise, we recently showed that intra-tumoral (IT)-injection of murine (M)CMV into poorly immunogenic B16 melanomas resulted in delayed tumor growth^51^. Finally, the Herbein lab demonstrated that human (H)CMV could inhibit the growth of hepatocellular carcinoma as well activate pathways involved in tumorigenesis^52,53^. Thus, it is unclear how CMV may affect the tumor environment and how these effects vary based on the type of tumor.

Given the disparate effects of CMV on various cancers, we wished to define the mechanism by which MCMV delayed tumor growth in the melanoma model. Importantly, the primary infected cells in the tumor were macrophages^51^, hinting that the virus was not oncolytic. Macrophages and other myeloid cells are important for the biology of CMV^54–60^, which utilizes these cells for systemic dissemination and as sites of latency. Thus, CMV has evolved mechanisms to recruit monocytes to the site of infection and alter their functional state. Additionally, tumor-associated macrophages (TAMs) have emerged as key regulators of the tumor microenvironment where they are predominantly anti-inflammatory or M2-like, and support tumor growth^61–63^. Recent work has shown that depleting these M2-like macrophages from the tumor or converting them to a more pro-inflammatory, or M1-like, state can induce tumor regression^64–67^. Thus, the goal of the present study was to define how MCMV modulated TAMs and whether the virus’ interaction with TAMs was important for delaying tumor growth in the melanoma model.

B16 melanomas are an immunologically “cold” tumor model, containing few infiltrating immune cells in the absence of interventions. Our data show that both wild-type and spread-defective MCMV induced tumor growth delay in a manner that depended on recruitment and infection of TAMs as well as the ability of the animals to mount adaptive immune responses. We found that the viral chemokine MCK2 was necessary to accumulate macrophages in the tumor after MCMV infection as well as to delay tumor growth. MCMV infection of M2-like macrophages drove them toward an M1-like state and MCMV infection of the tumor induced marked, but transient inflammation. However, repeated rounds of IT-MCMV could sustain and prolong the anti-tumor effect, suggesting that an active MCMV infection alters the tumor microenvironment in an immunologically “cold” melanoma model by actively recruiting and modulating TAMs, thereby promoting anti-tumor responses.

## Materials and Methods

### Mice

C57BL/6J or Rag KO (B6.129S7-*Rag1*^*tm1Mom*^/J) mice were purchased from The Jackson Laboratory and maintained in-house. For most studies, both male and female mice were used and were between 6 and 10 weeks at the time of tumor implantation or bone marrow harvest. The Institutional Animal Care and Use Committee at Thomas Jefferson University reviewed and approved all protocols.

### Tumor, Viruses and intratumoral injections

B16-F0 cells were purchased from ATCC, grown in DMEM with 1% penicillin-streptomycin and 10% FBS and frozen in a large batch of aliquots between passage 5 and 7. In all experiments, cells derived from this frozen batch were thawed, passaged once more, and implanted within seven days. Injected cells were always pigmented and the batch was certified negative for mycoplasma and other pathogens by IMPACT III testing on 10/30/2017 by IDEXX BioResearch. For tumor implantation, B16-F0s were resuspended in HBSS and injected subcutaneously in the shaved right flank of the animal as described previously^68^. Tumor growth was monitored by measuring length and width with a 6-inch digital caliper (Neiko). When tumors reached 20mm^2^ in area they were directly injected with MCMV or PBS as a control using an insulin syringe with 5×10^5^ pfu of virus in 30 µL of PBS, every other day for a total of three injections. WT-MCMV (K181) and spread-defective MCMV (ΔgL, based on the K181 backbone^10^), have been previously described. Viruses expressing an MCK2 with mutations in the chemokine domain (RM4511, hereafter called MCK2^mut^), or its matched wild-type counterpart (RM4503, hereafter called MCK2^WT^), as well as a virus completely lacking MCK2 (RM461, referred to as MCK2 KO) and its matched counterpart (RM427, referred to as MCK2 WT) were kindly provided by Dr. Edward Mocarski^69^. All viruses were grown as previously described^70^. After IT injections, tumors were monitored until they reached 100 mm^2^.

### Clodronate depletion

To deplete monocytic phagocytes, clodronate liposomes (Liposoma B.V., Amsterdam) were diluted in PBS and injected at a concentration of 10 mg/kg IP and 5 mg/kg IT as previously described^64^. Liposomes were injected beginning 1 day before intratumoral injections (e.g. day -1) and were repeated on days 2, 4, 7 and 11 or until sacrifice if tumors reached 100mm^2^ before day 11. The amount of liposomes was adjusted weekly based on animal weight.

### Immunofluorescence microscopy

Tumors were rapidly frozen in O.C.T. compound and cut tumors into 7 µM sections by the Thomas Jefferson University Pathology Core. Sections were placed in cold acetone for 10 minutes, rehydrated with Tris-buffered saline (TBS) for 20 minutes, and blocked with blocking buffer (TBS + 3% BSA and 0.1% Tween-20) for 20 minutes. In all figures, sections were stained with antibodies specific for F4/80 (clone BM8), CD11b (clone M1/70), and DAPI in blocking buffer for one hour. Samples were imaged using the Nikon A1R fluorescent confocal microscope. Macrophage number per mm^2^ image was calculated from the count of F4/80+, CD11b+ macrophages per image from 6 images per tumor and 3-4 tumors per treatment.

### Bone marrow culture and in vitro viral infections

L929-conditioned media for growing macrophages was produced by plating 7.2×10^5^ L929 cells in T-150 flasks. Supernatant was harvested after 7 days, replaced with fresh media and harvested again after 14 days. All L929 conditioned media was combined and frozen for later use. The bone marrow from femurs and tibias of C57BL/6J mice was flushed and incubated in bone marrow macrophage media consisting of DMEM with 10% L929-conditioned media, 10% fetal bovine serum, and 1% penicillin-streptomycin. Non-tissue culture treated petri dishes were used to facilitate macrophage recovery from the plastic. On day 7 of the culture, macrophages were re-plated at 6×10^5^ cells/well in a 6 well plate. M2-like macrophages were polarized using 40 ng/mL IL-4 on day 8. On day 9, MCMV was added at a multiplicity of infection (MOI) of 5 to half of the IL-4-treated wells without removing the polarizing media. For comparison, M1-like macrophages were polarized by adding 20 ng/mL of IFN-γ and 1 µg/mL LPS on day 9 of culture in parallel wells. 24 hours after infection, all macrophages were harvested for analysis.

To test the ability of MCK2^mut^ MCMV and its wild-type counterpart to infect macrophages (Figure 4E), bone marrow derived macrophages (BMDMs) were produced as above and infected with either virus at an MOI of 1 on the 8th day of culture (with no cytokines added for polarization). After 24 hours, the cells were fixed and permeabilized with a 1× dilution of True-Nuclear Transcription Factor kit (BioLegend) following the manufacturer’s instructions. Cells were blocked overnight in PBS with 0.2% BSA, 0.1% gelatin, and 0.1% Triton X-100 and then stained for expression of MCMV pp89 using clone 6/58/1^71^ for 1 hour, followed by anti-mouse IgG2b (clone RMG2b-1, BioLegend). Infected cells were identified using a Nikon eclipse Ti confocal microscope and analyzed with ImageJ (https://fiji.sc/).

### Flow cytometry

Tumors were placed in digestion media (HBSS with 0.5 mg/mL collagenase type I and 60 U/mL DNAse,), minced using the gentleMACS Octo Dissociator (Miltenyi Biotec) on m_lung_01_01 and m_lung_02_01 settings, and incubated at 37°C with rotation for 30 minutes. Following the incubation, cells were separated with Lymphoprep (Stem Cell Technologies), following the manufacturer’s instructions and stained for flow cytometry. Antibodies specific for CD11b (clone M1/70), F4/80 (clone BM8), I-A/I-E (clone M5/114.15.12), CD11c (clone N418), CD206 (clone C068C2), CD45.2 (clone 104), PD-L1 (clone 10F.9G2), and Ly6C (clone HK1.4), all from Biolegend, were used to define the phenotype of tumor infiltrating macrophages. CountBright Absolute Counting Beads (Biolegend) were also included. Lymphocytes were excluded by dumping cells staining positive for CD3 (clone 17A2), CD19 (clone 1D3/CD19), and NK1.1 (clone PK136). Representative gating strategies are shown in Supplemental Figure 1A-B. For RPPA validation, BMDMs (Supplemental Figure 1B-C) were gated on CD11b and F4/80 and co-stained with anti-phospho-S6 S235/S236 (clone cupk43k from eBioscience). All FACS data was collected on a BD LSRFortessa or BD FACSCelesta and analyzed using FlowJo software v10.4.2 (TreeStar).

### RPPA analysis

BMDMs (polarized to M1 and M2, or unpolarized M0) were infected with MCMV at an MOI of 5 or left uninfected. After 24 hours cells were rinsed with PBS and lysed for 20 minutes with RPPA lysis buffer (1% Triton X-100, 50mM HEPES, 150mM NaCl, 1.5mM MgCl_2_, 1mM EGTA, 100mM NaF, 10mM Na pyrophosphate, 1mM Na_3_VO_4_, 10% glycerol). Lysed cells were scraped and pelleted. The protein concentration in the supernatant was analyzed using a Bradford Assay and adjusted to 1 mg/mL. Samples were mixed with SDS and denatured by boiling before being sent for analysis. Data were normalized by protein loading levels, as described at https://www.mdanderson.org/research/research-resources/core-facilities/functional-proteomics-rppa-core/faq.html. The limma package (v3.36.2)^72^ was used to perform differential expression analysis. Cell lineage trajectory analysis was done using the monocle package (v2.8.0)^73–75^. Heatmaps were generated using the pheatmap package (v1.0.10 https://CRAN.R-project.org/package=pheatmap). Scatter plots were generated using the ggplot2 package (v3.0.0 https://ggplot2.tidyverse.org/). Data analyses were performed using R (v3.5.1 http://www.R-project.org/).

### Quantitative PCR

RNA was extracted from infected and uninfected BMDMs (Figure 5A) or from whole tumor homogenates (Figure 6A) produced by pushing tumors through a 70 µm filter. RNA was isolated using the RNeasy Mini Kit (QIAGEN) and cDNA was produced using a high capacity cDNA reverse transcription kit (Applied Biosystems). For all experiments, transcripts were detected using individually designed primers (Integrated DNA technologies) and probes for cytokines (Integrated DNA technologies) or MCMV E1 (applied biosystems). A predesigned qPCR assay (Integrated DNA technologies) was used to detect the housekeeping gene beta actin and all samples were run on the StepOnePlus system (Applied Biosystems). In macrophage studies, transcripts were normalized to beta actin in each sample and compared to uninfected and unpolarized (M0) macrophages to obtain a ΔΔCT value. Data are expressed as fold change over M0 macrophages using 2^−(ΔΔCT)^. For tumor homogenates, transcript concentrations were normalized to beta actin in each sample and recorded using the formula 2^−(ΔCT)^.

### Statistical analysis

Statistics were performed using GraphPad Prism v6 or R (v3.3.1 R-project.org). If normally distributed (as assessed by the Kolmogorov-Smirnov test), data were analyzed using a two-tailed t test. If data were non-normally distributed, a Mann-Whitney test was performed instead. For tumor doubling times, the tumor measurements were log transformed and a linear regression was used to model the rate of tumor growth as a function of time. Finally, a log-rank (Mantel-Cox) test was used to compare Kaplan-Meier survival curves.

### Data Availability

All data is freely available upon request. RPPA data can be found in Supplemental Table 1.

## Results

### Intratumoral injections of WT-MCMV lead to delayed tumor growth

We have previously reported that IT injections of MCMV lead to delayed growth of established B16-F0 melanomas. In this treatment schedule, MCMV was injected directly into the melanoma lesions every other day for a total of three injections beginning once the tumors reached approximately 20mm^2^ in area (Figure 1A)^76^. Consistent with our previous data, injections of WT-MCMV delayed tumor growth and significantly prolonged the time for tumors to reach the pre-determined end point of 100mm^2^ (Figure 1B-C). Median survival after reaching 20 mm^2^ was 9 days for the mice treated with IT-PBS and 19.5 days for mice treated with IT-MCMV. Tumor doubling-time was also assessed by linear regression of log-transformed growth data. This is a less sensitive measure of tumor growth since it includes data points from the exponential growth phase after treatment and excludes animals that clear their tumors. Nevertheless, tumor doubling time was significantly increased by IT-MCMV injections (Figure 1D). Importantly, nearly identical results were obtained with a spread-defective version of MCMV (Figure 1B-D), which lacks the essential glycoprotein gL and can only produce non-infectious viral particles in non-complementing cells^10^. These data confirm our previous results that MCMV, without the inclusion of any tumor-derived vaccine antigens, can cause significantly delay tumor growth when injected intratumorally. Moreover, these data strongly argue that MCMV is not acting as an oncolytic virus in this model since even a spread-defective version of MCMV was similarly able to delay tumor growth.

**Figure 1.**
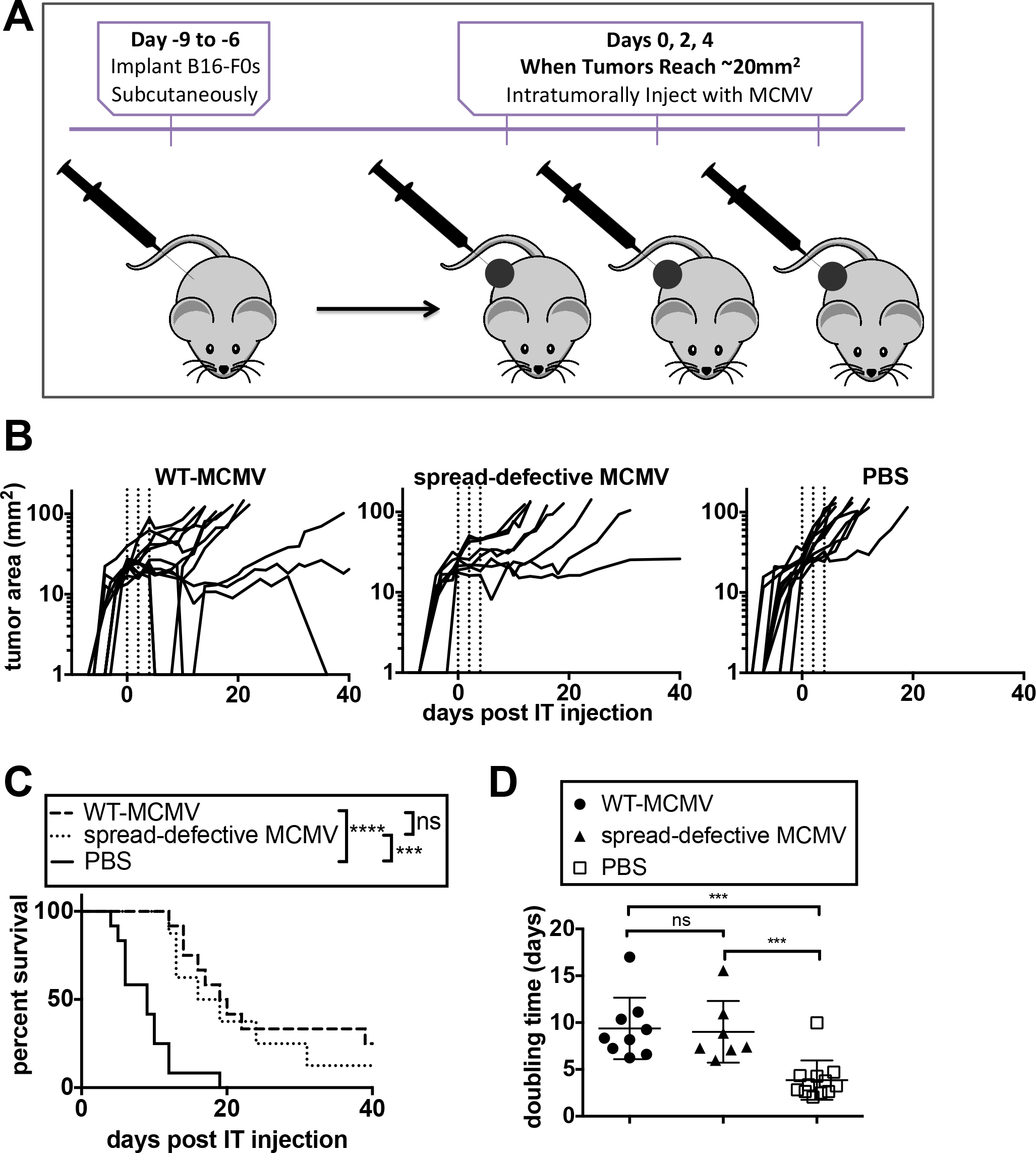
Intratumoral injections of WT-MCMV lead to delayed tumor growth **A)** Schematic showing the schedule for intratumoral injections. B16-F0s were implanted subcutaneously in the right shaved flank of the animal. Once the tumor reached around 20mm^2^, MCMV was injected into the tumor every other day for a total of three injections corresponding to day 0, day 2, and day 4. **B)** C57BL/6J mice were injected IT with WT-MCMV (n=12), spread-defective MCMV (n=8), or PBS (n=12). Tracings are on a logarithmic scale and show tumor growth in individual animals. Dotted lines indicate the days of IT injection. The length and width of the tumor were measured until reaching a size of 100mm^2^. **C)** Kaplan-Meier survival curves were plotted using the day when the tumors reached 100mm^2^ as the endpoint. Data were compared with a log-rank test. **D)** Tumor doubling times for each animal were calculated from day 0 (start of treatment) to sacrifice, excluding mice that cleared their tumor.

### Monocytic phagocytes are necessary but not sufficient for tumor growth delay

Our previous data showed that MCMV preferentially infected TAMs in this model^76^. In order to determine the importance of TAMs, we depleted phagocytic cells with clodronate-filled liposomes delivered IP and IT as described in the materials and methods. Depletion of phagocytes had no effect on the growth of tumors treated with IT-PBS (Figure 2A-C). Moreover, IT-MCMV still delayed tumor growth in mice treated with control liposomes (Figure 2A-C), although tumor growth was slightly accelerated compared to tumors that did not receive any liposomes (median survival of 15.5 days as compared to 19.5 days in Figure 1B-C). Strikingly however, clodronate-filled liposomes abolished the ability of IT-MCMV to delay tumor growth, prolong survival or increase doubling times (Figure 2A-C). The successful depletion of macrophages (F4/80+, CD11b+) from the tumor environment was confirmed by histology one day after the final IT injection (Fig. 2D). Therefore, the data indicate that monocytic phagocytes are necessary for the anti-tumor effect of IT-MCMV.

**Figure 2.**
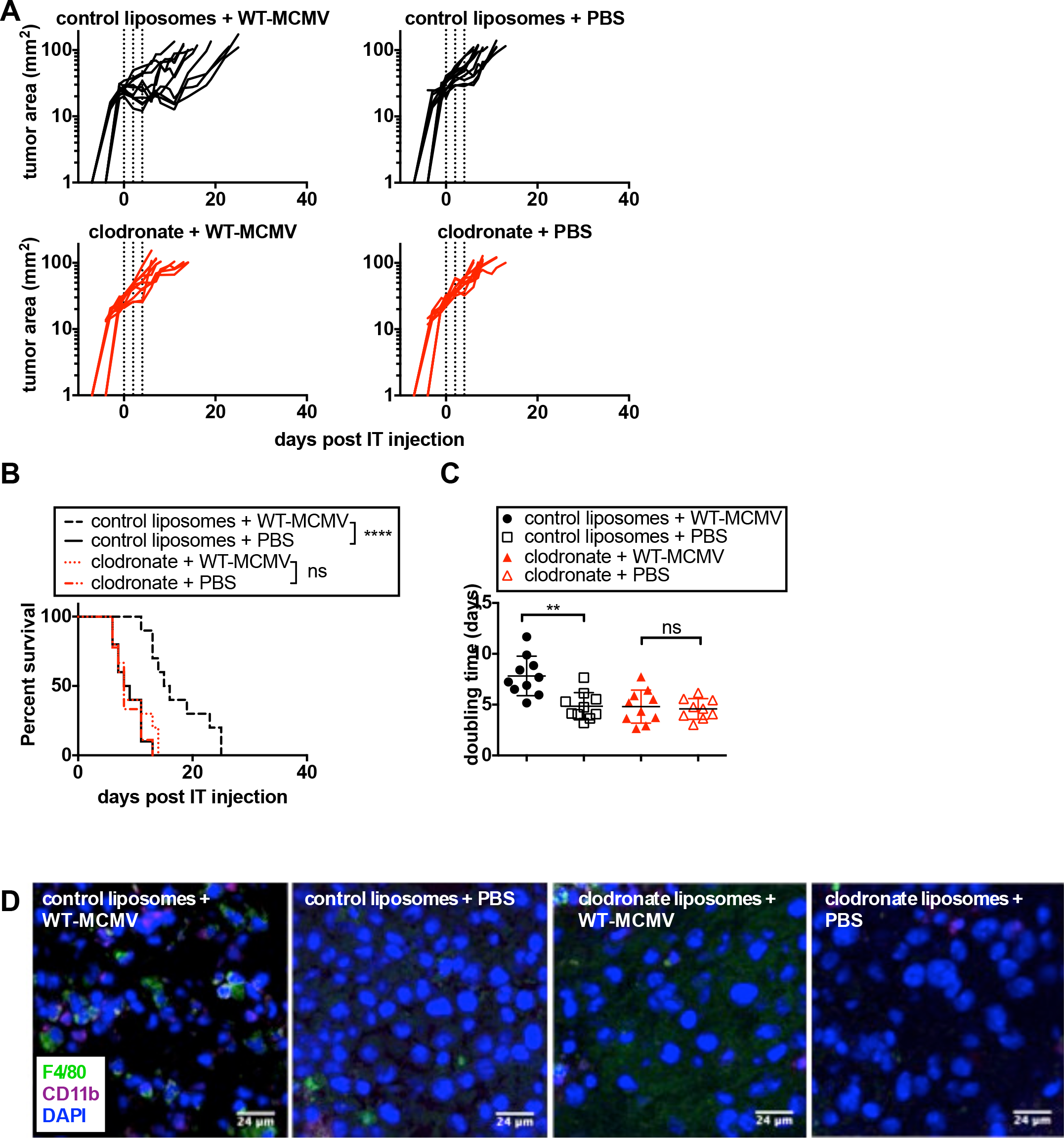
Monocytic phagocytes are necessary for tumor growth delay. Mice received control liposomes and WT-MCMV (n=10), control liposomes and PBS (n=10), clodronate liposomes and WT-MCMV (n=10), or clodronate liposomes and PBS (n=10). Tracings of tumor growth (**A**), animal survival (**B**), and tumor doubling times (**C**) are shown as in Figure 1. **D)** Representative images show the relative amount of F480+CD11b+ macrophages in each group after treatment.

We previously showed that antibody depletion of CD8+ cells, which includes both T cells and dendritic cells, ablated the efficacy of IT-MCMV^76^. Since our new data show that clodronate depletion of phagocytes prevented MCMV from delaying tumor growth, it remained possible that anti-CD8 antibodies accelerated tumor growth by depleting CD8+ dendritic cells. Thus, to confirm that adaptive immune cells are required for tumor growth delay, we tested the efficacy of IT injections of spread-defective MCMV in Rag KO animals. Interestingly, treatment had a small, but significant effect on survival and tumor doubling times (Figure 3A-C). Nevertheless, efficacy was severely blunted in these Rag KO mice, with a median survival of only 13 days (Figure 3B). These data confirm that adaptive immune mechanisms are required for anti-tumor immunity in addition to monocytic phagocytes.

**Figure 3.**
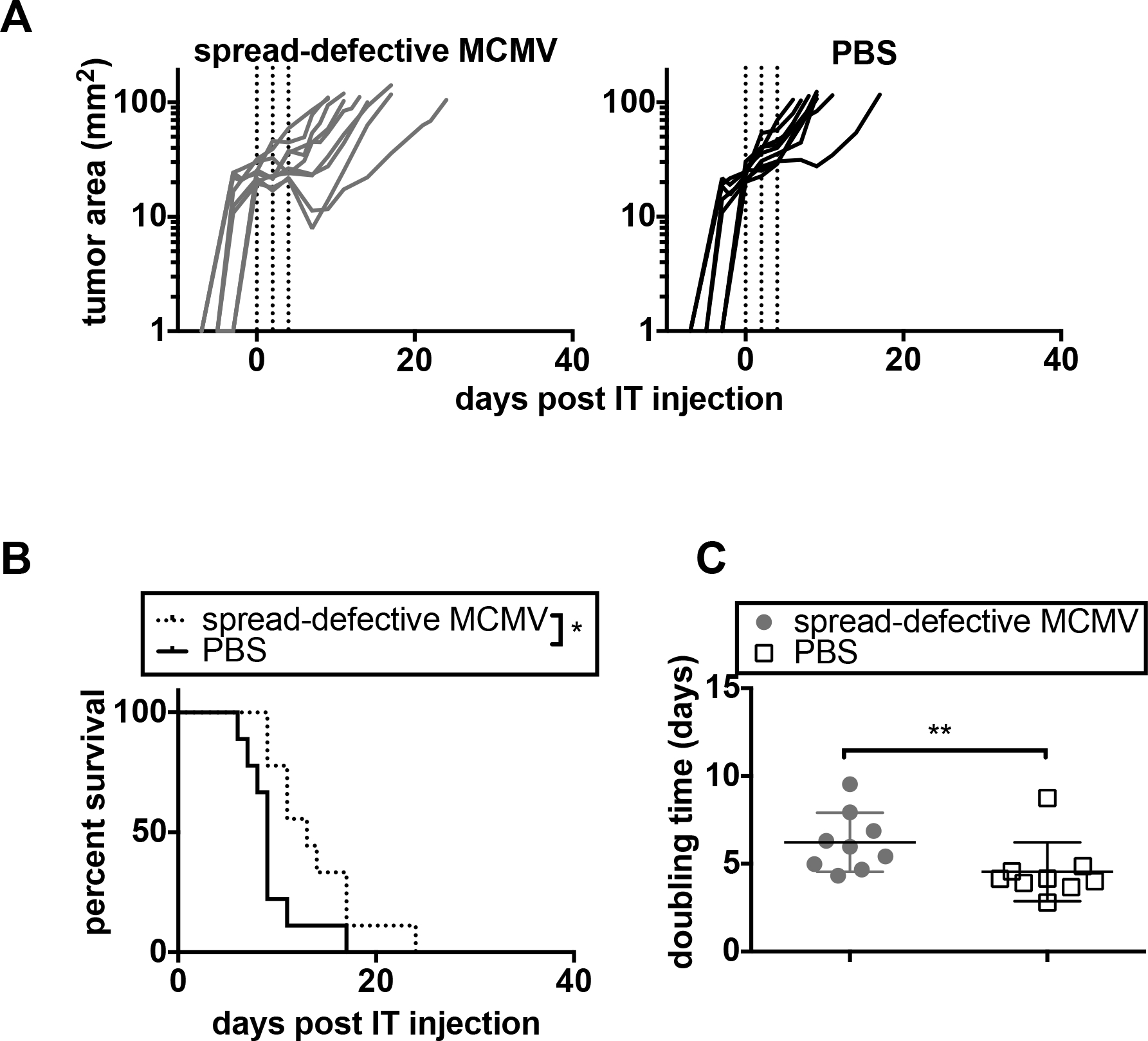
IT-MCMV is ineffective in Rag-knockout mice. Rag KO mice were injected IT with spread-defective MCMV (n=9) or PBS (n=9) every other day for a total of three injections. Tracings of tumor growth (**A**), animal survival (**B**), and tumor doubling times (**C**) are shown as in Figure 1.

### MCK2 mediated recruitment of new macrophages is indispensable for tumor growth delay

The MCMV encoded chemokine MCK2 has been implicated in the recruitment of monocytes to the site of infection^77–79^. Thus, we hypothesized that MCK2 might be a critical regulator of MCMV driven anti-tumor immunity. To test this, we used a mutant MCMV in which the dual cysteines of the chemokine domain of MCK2 were mutated (MCK2^mut^) along with a matched wild-type strain (hereafter called MCK2^WT^)^69^. Remarkably, the MCK2^mut^ MCMV was significantly less effective at delaying tumor growth or increasing survival when compared to the MCK2^WT^ virus (Figure 4A-C). IT treatment with MCK2^mut^ MCMV still increased animal survival and tumor doubling times compared to PBS-treated tumors, although the effects were less dramatic. MCK2 has been implicated in the entry of MCMV into macrophages^79^. However, the MCK2^mut^ virus was able to infect macrophages equally to its wild-type counterpart (Figure 4E). Therefore, these data strongly suggest that MCK2 was required to recruit macrophages into the tumor. Importantly, we repeated this experiment with a second, independently generated mutant MCMV that completely lacked MCK2 (MCK2 KO) compared to its matched wild-type control virus. This virus completely lacking MCK2 exhibited an even more striking loss of therapeutic efficacy, with no increase in survival or doubling time compared to PBS-treated tumors (Figure 4F-H)^60,78^. These data show that wild-type viral MCK2 is needed for the anti-tumor effect of IT-MCMV.

**Figure 4.**
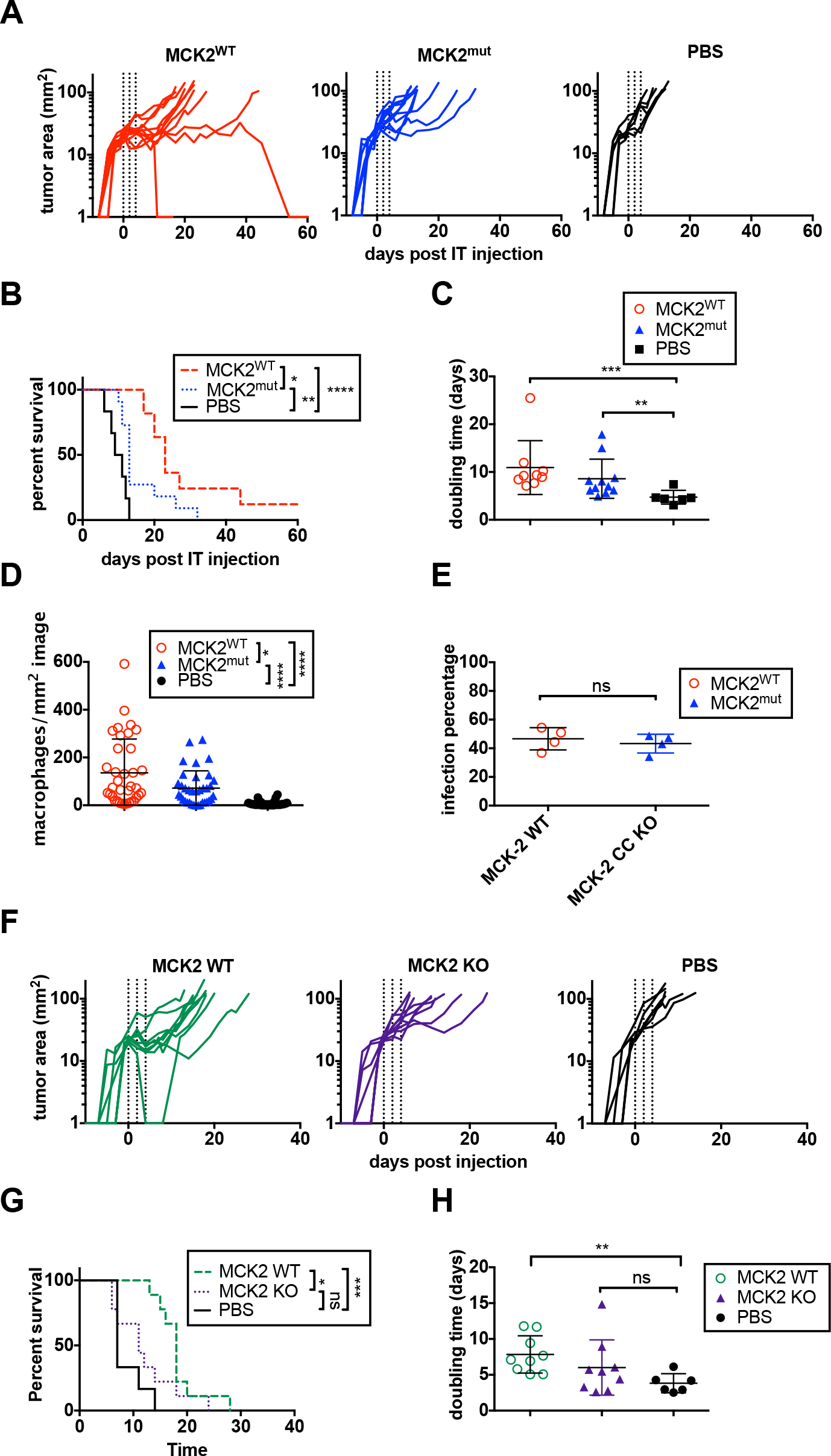
MCK2 mediated recruitment of macrophages is indispensable for tumor growth delay. Tumor-bearing mice were treated IT with MCK2^mut^ virus (n=11), MCK2^WT^ virus (n=11), or PBS (n=7). Tracings of tumor growth (**A**), animal survival (**B**), and tumor doubling times (**C**) are shown as in Figure 1. **D)** Macrophage numbers were measured by immunofluorescent histology for the presence of F4/80+, CD11b+ cells in 6 representative images of tumors treated with MCK2^mut^ virus (n=4 tumors), MCK2^WT^ virus (n=4), and PBS (n=3). Shown is the density of macrophages per mm^2^ of each analyzed image. **E)** The frequency of BMDM that were infected by MCK2^WT^ or MCK2^mut^ viruses is shown. Four individual wells of macrophages were assessed and the frequency of infected cells averaged from 3 images from each well. A two-tailed t-test was used to test for differences. Tumor-bearing mice were also treated IT with MCK2 KO virus (n=9), MCK2 WT virus (n=9), or PBS (n=6) to confirm initial findings with an independently generated virus. Tracings of tumor growth (**F**), animal survival (**G**), and tumor doubling times (**H**) are shown.

Significantly fewer macrophages were detected in the tumors injected with MCK2^mut^ virus compared to MCK2^WT^ MCMV (Figure 4D), although both MCK2^WT^ and MCK2^mut^ viruses increased the number of TAMs compared to PBS-injected tumors. Thus, the loss of the MCK2 chemokine impaired the recruitment of new monocytes to the tumor. Additionally, we noted that macrophages seemed to infiltrate into the center of tumors treated with the MCK2^WT^ virus compared to MCK2^mut^ virus (Supplemental Figure 2), suggesting that MCK2 may impact both the number and localization of TAMs after IT infection.

### MCMV infection re-polarizes macrophages to become more inflammatory

In order to better understand the effect MCMV infection was having on the recruited macrophages, we produced bone marrow derived macrophages (BMDMs) and either left them unpolarized (M0), or polarized them to an M2-like or an M1-like state as described in the materials and methods. After polarization, each sub-type was infected with MCMV and we assessed changes in the expression of various transcripts relative to uninfected M0 macrophages. The data show a trend of MCMV infection driving cells towards a more M1-like state (increased iNOS, increased TNF-α, reduced arginase), with changes in iNOS reaching statistical significance (Figure 5A).

**Figure 5.**
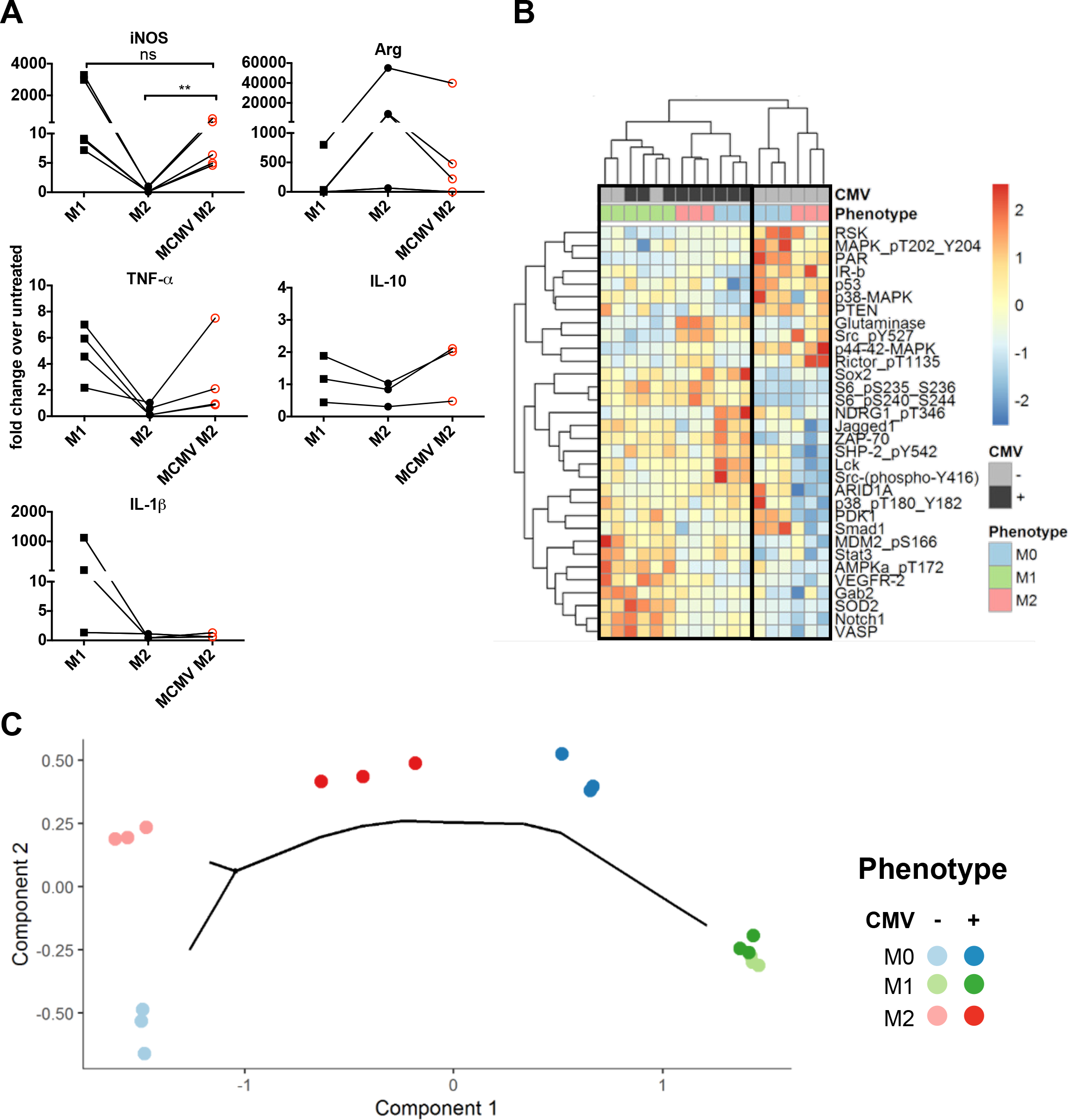
MCMV infection re-polarizes macrophages to become more M1-like **A)** Polarized and unpolarized BMDMs were infected with MCMV or mock-infected and assessed by qPCR for iNOS (n=5 independent macrophage cultures), TNF-α (n=4), IL-1β (n=3), Arg (n=4), IL-10 (n=3). Fold change of each transcript relative to untreated M0 macrophages is shown using the calculation 2^−(ΔΔCT)^, and data were compared using the Mann-Whitney test. **B)** Unsupervised hierarchical clustering of RPPA analyses of infected and uninfected macrophages that were polarized or unpolarized. Shown is a heatmap of antibodies that showed significant differences (BHFDR < 0.05) with or without MCMV infection or between the M1 and M2 polarized states. **C)** Monocle pseudo-time plot of RPPA data assessing macrophage cell differentiation. Starting from time point 0 (bottom left corner), two distinct lineages appear, separating uninfected M1 and M2 macrophages (M1 to the right and M2 to the top left). In contrast, both MCMV-infected M0 and MCMV-infected M2 macrophages appear on the M1 lineage branch. MCMV-infected M1 macrophages appear at the end of the M1 branch, meaning that there are no differences between the M1 and M1 + MCMV samples.

For a more comprehensive analysis, we used reverse phase protein array (RPPA), a quantitative proteomics approach. For this, BMDMs were cultured and infected as above, and cell lysates were subjected to RPPA analysis. Unsupervised clustering of protein quantitation shows that M1-like macrophages are distinct from both M0 and M2-like macrophages (Figure 5B). However, infection of both M0 and M2 macrophages led them to be clustered with M1 macrophages and with MCMV-infected M1 macrophages. We also used a Monocle pseudotime plot to define trajectories of differentiation between each cell population. As shown in Figure 5C, M0, M1 and M2 macrophages represent distinct branches of differentiation prior to infection. Infection by MCMV of both M0 and M2 macrophages drove them onto the M1 branch. In contrast, uninfected and infected M1 macrophages were virtually indistinguishable. Thus, MCMV infection can polarize macrophages to become more M1-like.

To validate the RPPA data, we chose to assess the phosphorylation of S6, which was strongly increased by MCMV infection in the RPPA analysis (Figure 5B). We could clearly detect an increase in S6 phosphorylation by flow cytometry after MCMV infection (Supplemental Figure 1C), which is consistent with previous work using human (H)CMV^80^. Overall, these data indicate that MCMV has the ability to re-polarize M2-like macrophages toward a more inflammatory state.

### The effect of IT-MCMV is transient, but repeated IT injections recapitulate the therapeutic response

The effect of IT-MCMV is transient and tumors began to regrow in most mice approximately 7-10 days after therapy began. We harvested tumors on days 5, 8 and 11 after the start of IT therapy and found that expression levels of iNOS, TNF-α, IL-1β, arginase and IL-10 were all increased by day 5 after the first IT-MCMV injection (Figure 6A). Inflammatory transcripts (iNOS, TNF-α, IL-1β) also seemed to be sustained at higher levels on Day 8, but did not reach significance over PBS. The specific contribution of TAMs to these transcripts could not be directly assessed because baseline levels of these transcripts were increased by clodronate liposomes alone (not shown). Regardless, these changes were transient: by day 11, all transcripts had returned to the baseline levels seen in PBS-treated tumors. This time-point correlates with a loss of viral transcription in the tumor (Figure 6B) and approximately aligns with the beginning of tumor regrowth (e.g. Figure 1). Thus, the inflammatory changes associated with a therapeutic effect were not sustained as the virus was controlled. Additionally, we recovered TAMs at all 3 time points and assessed their phenotype by flow cytometry (Figure 6C). Therapy with IT-MCMV was associated with the emergence of a distinct F4/80^hi^Ly6C^int^ macrophage population on day 8. These cells expressed high levels of PD-L1 and MHC-II, indicating their activation (Figure 6D). However, this population was no longer evident by day 11. Together, these data indicate that a single round of IT-MCMV therapy transiently alters the tumor microenvironment by recruiting and activating TAMs and altering the inflammatory milieu.

**Figure 6.**
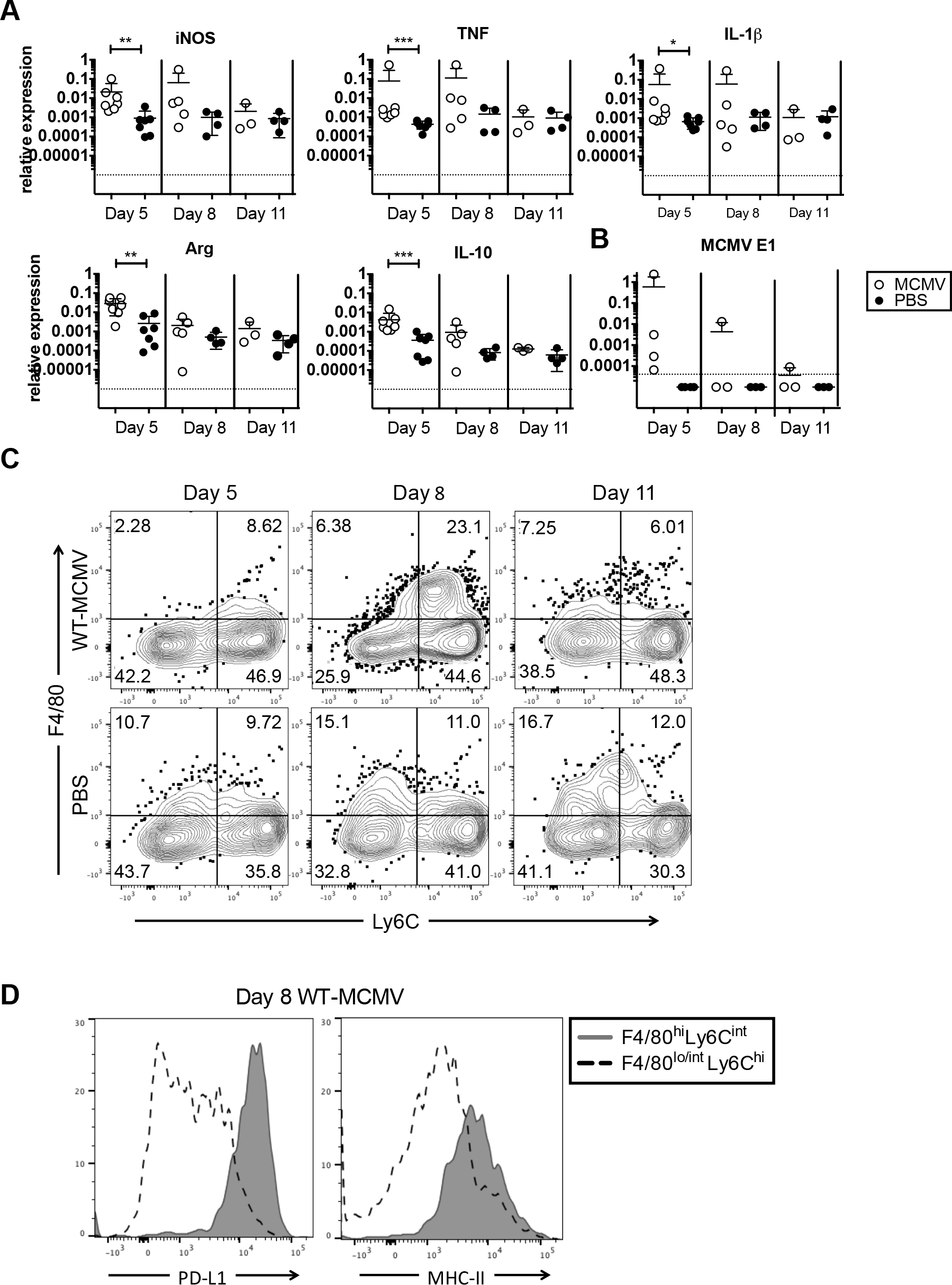
Macrophage-associated transcript and resident macrophage populations are transiently increase following IT-MCMV injections **A)** Relative expression of the indicated transcripts from whole tumor homogenate is shown. Tumors treated with WT-MCMV or PBS were harvested on day 5 (WT-MCMV and PBS n=7), day 8 (WT-MCMV n=5, PBS n=4), and day 11 (WT-MCMV n=3, PBS n=4) after the initial IT injection. Data were compared using the Mann-Whitney test. **B)** Relative expression of viral transcript is shown as is A. Tumors treated with WT-MCMV or PBS were harvested on day 5 (WT-MCMV and PBS n=4), day 8 (WT-MCMV and PBS n=3), and day 11 (WT-MCMV and PBS n=3) after the initial IT injection. **C)** Concatenated FACS plots show F4/80+, Ly6C+ macrophages on days 5, 8, and 11 after IT-MCMV or IT-PBS. Plots are concatenated from four samples each and represent populations previously gated on CD45+, NK1.1-, CD3-, CD19-, and CD11b+. **D)** Histograms show PD-L1 and MHC-II expression by the indicated macrophage populations from IT-MCMV-treated tumors shown in Figure B, day 8.

Since the effects of IT-MCMV were transient and tumor re-growth coincided with a loss of viral activity in the tumor, we wondered whether additional IT-MCMV treatment would delay tumor growth further. To test this, we treated tumors with IT-MCMV as above (Figure 1A) when the therapy was no longer working as indicated by multiple days of positive growth after the period of stable disease. When treated tumors re-surpassed a size of 20mm^2^, the animals received 3 additional injections of MCMV (denoted as 6 WT-MCMV) or 3 injections of PBS (denoted as 3 WT-MCMV + 3 PBS). This second round of IT-MCMV therapy further delayed tumor growth and significantly improved overall survival (Figure 7A-B). Tumor doubling times were also significantly increased to 19.2 days for animals that received 6 doses of IT-MCMV versus 12.2 days for 3 doses of MCMV plus 3 doses of PBS (p=0.046, not shown). Remarkably, when the time required for tumors to reach 100mm^2^ was normalized to the days after the last MCMV treatment (the 3rd or the 6th overall injection), there was no significant difference (Figure 7C), indicating that the second round of therapy increased overall survival by recapitulating the original effect. Interestingly, tumors taken one day after the 6^th^ injection with WT-MCMV or one day after the last injection in the 3 WT-MCMV +3 PBS group showed similar numbers of macrophages in the tissue (Figure 7D-E). These data, combined with the near absence of TAMs in tumors that never received MCMV (Figure 4D) suggest the TAMs persisted from the initial round of IT-MCMV treatment. Thus, loss of TAMs cannot explain the tumor progression. Rather, our data suggest that active viral infection was needed to delay tumor growth and that repeated IT injections may be able to sustain the inflammatory environment and the M1-like polarization state of TAMs. Collectively, our data show that IT-MCMV infection recruits monocytic phagocytes to the tumor via the viral chemokine MCK2 to delay tumor growth, and suggest that infection of these TAMs by MCMV polarizes them to an M1-like state that is transient, but can be sustained by repeated IT-MCMV administrations to promote functional anti-tumor immune responses.

**Figure 7.**
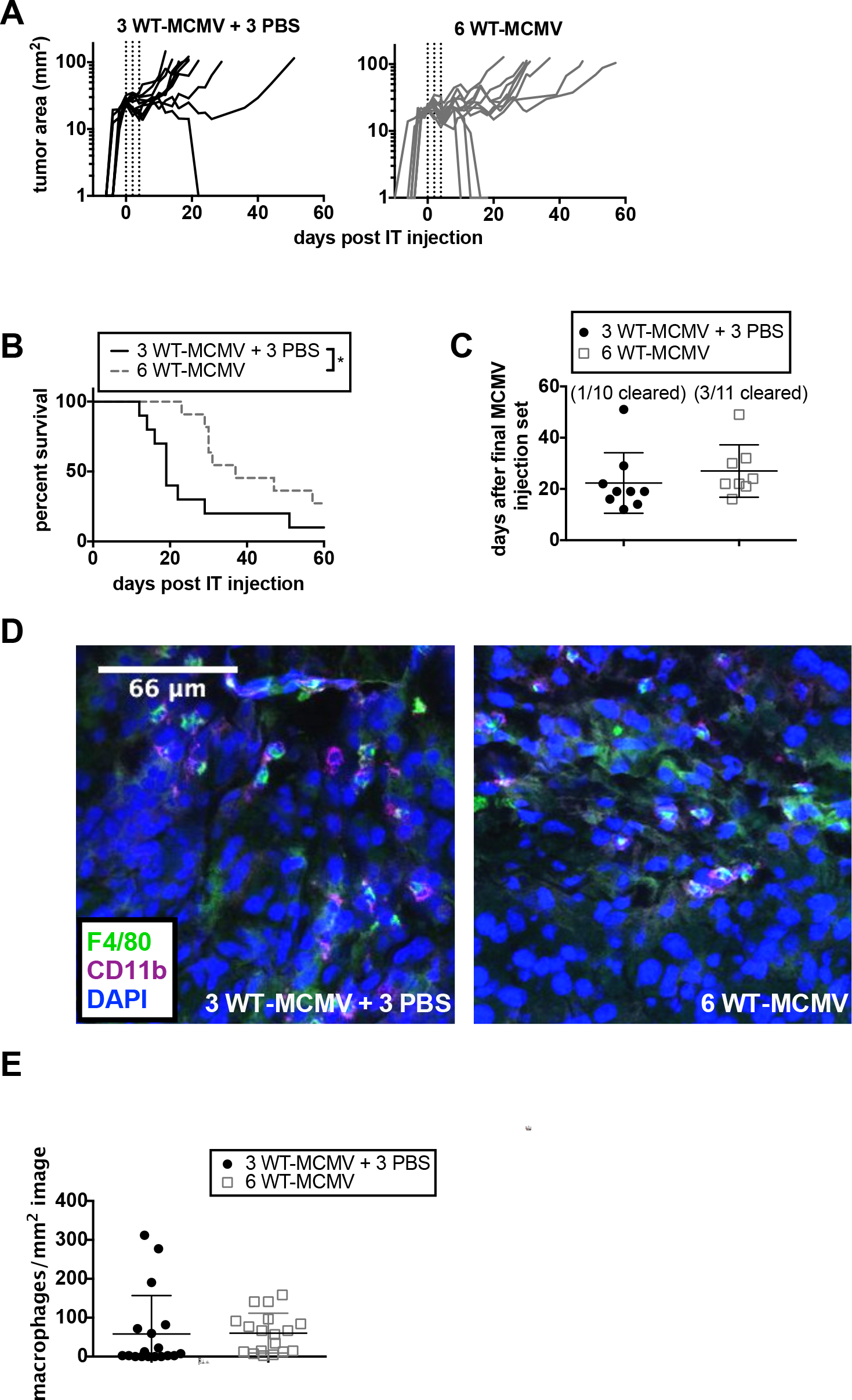
Multiple tumor injections recapitulate the original effect of IT-MCMCV therapy and cause a significant increase in survival. Tumor-bearing mice were treated IT with 3 injections of WT-MCMV followed by 3 injections of PBS (n=10) or 3 more injections of WT-MCMV (n=11). The second round of treatment began when tumors surpassed 20mm^2^ once again. Tracings of tumor growth (**A**) and animal survival (**B**) are shown as in Figure 1. **C)** The time to sacrifice for each treatment is shown normalized to the last set of MCMV injections. Using immunofluorescent histology, representative images from the day after the sixth injection show the number of macrophages in the tumor which is further quantified (**E)** by counting the number of F4/80+CD11b+ cells in 6 images from tumors treated with 6 injections of WT-MCMV (n=3 tumors) or 3 injections of WT-MCMV followed by 3 injections of PBS (n=3). Shown is the density of macrophages per mm^2^ of each analyzed image. Data were compared using a two-tailed t-test, but there was no significant difference.

## Discussion

It is increasingly clear that viral modulation of the tumor microenvironment is a promising strategy for tumor immune therapy. CMV can effectively be used as a vaccine platform to generate anti-tumor immune responses in multiple models^16–22^. However, the ability of CMV to infect and establish latency in myeloid cells means that any tissue or tumor that recruits monocytes will likely recruit CMV-infected monocytes. Since CMV can be found in a variety of solid tumors^33,34^ and it has been hypothesized that CMV may promote or exacerbate tumor growth, it is important to discern the impact of CMV on tumor microenvironments. We had previously observed that MCMV could delay the growth of the poorly immunogenic B16 melanoma and the current study aimed to define the mechanism behind this unexpected outcome. Our data suggest that viral infection inhibits tumor growth by actively recruiting monocytes that become TAMs. TAMs were subsequently infected and activated by infection. Therefore, we propose that MCMV engages adaptive immune responses in the tumor by recruiting and activating TAMs. However, once viral replication was lost, tumors began to regrow. Tumors in most mice returned to an exponential growth phase beginning 7 to 10 days after the initiation of therapy, which corresponded with a loss of viral activity in the tumor (Figure 6B). We found that macrophage numbers remained elevated as tumors began to regrow and that a second round of MCMV infection was able to re-delay tumor growth and increase the number of mice that cleared their tumors (Figure 7). These data suggest that active MCMV infection may be the key for driving immunity that can delay tumor growth.

This study is limited to the anti-tumor effects of MCMV during an active intratumoral infection in a model using tumor cells that are already transformed and are poorly infected by the virus. Thus, our data do not speak to the potential of MCMV to promote transformation of normal cells nor do they suggest how latency may lead to oncomodulation. However, our data do indicate that virus-recruited macrophages remained in the tumor. In settings where these macrophages can be converted to an anti-inflammatory M2-like state by the tumor, it is possible that the recruited macrophages could ultimately support tumor growth. In our hands however, when the tumor started to regrow after IT-MCMV, the tumor doubling time was still reduced (i.e. growth was still slower) when compared mice that never received MCMV (data not shown). Moreover, we have seen slower growth of untreated tumors in mice latently infected with MCMV (data not shown). These data could indicate that latent virus may not be detrimental in some cases, although further research is needed to fully define the effects of MCMV latency on tumor growth and progression.

Viral recruitment of TAMs in this model was dependent on the MCK2 chemokine (Figure 4, Supplemental Figure 2). Human CMV expresses the chemokine UL128^81^, which is the functional homologue of MCK2. Although MCK2 is known to attract CX3CR1+ monocytes to sites of infection^77^, the specific chemokine receptor as well as the functional state of the recruited monocytes remains undefined. Thus, future work will be needed to determine which monocytes are specifically recruited to the tumor and whether these are similar to monocytes recruited by this or other tumors under various conditions.

Our data also suggest that MCMV induced repolarization of TAMs to a more M1-like state. Similarly, HCMV can drive monocytes and macrophages to become M1-like^44,82^ and it will be interesting to learn whether human and mouse macrophages are similarly affected by viral infection and which molecular pathways are important for inducing a therapeutic effect. Likewise, future studies will need to determine whether viral infection of TAMs *per se* is needed to promote the M1-like state, or whether activation of pattern-recognition receptors on newly recruited TAMs is sufficient to prolong survival.

Our data suggest that CMV may offer a unique tool to directly target macrophages in the tumor environment through both recruitment and infection. Activation of TAMs has strong potential to provide synergy with current immune therapy treatment strategies, especially immune checkpoint blockades. Most people fail to respond to immune checkpoint therapies in isolation^83^ and immense effort is being devoted to rationally design combinations of therapies that can increase the number of responding patients. One of the main differences between responders and non-responders may be the degree of immune infiltration into the tumor. Thus, the B16-F0 tumor model is appropriate to test the virus in this context as it is very poorly immunogenic and checkpoint blockade therapies are mostly ineffective^76,85^. Our data suggest that MCMV was not acting as an oncolytic virus as it did not have to spread from the infected cells to delay tumor growth (Figure 1). Moreover, tumor growth delay was markedly reduced in the absence of an adaptive immune responses (Figure 3), suggesting that immune activation was an important part of MCMV’s impact on tumors. Emerging evidence suggests that immune activation in the tumor environment is an important feature of both oncolytic and non-oncolytic pathogens used for therapy, and may improve responses to immune checkpoint therapies^84^. Although the usefulness of MCMV infection as a combination therapy was not the goal of this study, we have previously shown that the combination of IT-MCMV with PD-L1 blockade resulted in clearance of ~70% of established tumors^76^. Thus, recruitment and modulation of tumor-associated monocytic phagocytes by MCMV may potentiate the efficacy of checkpoint blockades in immunologically “cold” tumor environments. Defining the mechanisms supporting this synergy will be a major goal of future work.

Overall, our study demonstrates that WT-MCMV is effective at generating anti-tumor immunity during periods of active viral replication. IT-MCMV effectively alters the tumor microenvironment by recruiting and polarizing macrophages in the tumor and engaging adaptive immune responses to delay tumor growth in a model resistant to current immunotherapies.

## Acknowledgments

This work was supported by a grant from the American Cancer Society awarded to C.M.S. and the Dr. Miriam and Sheldon G. Adelson Medical Research Foundation to A.E.A. The RPPA studies were performed at the Functional Proteomics Core Facility at The University of Texas MD Anderson Cancer Center, which is supported by the NCI Cancer Center Support Grant (CA16672). Immunofluorescent images were captured using the SKCC bioimaging shared resource (NCI 5 P30 CA-56036) with assistance from Maria Yolanda Covarrubias. Additional microscopy support was provided by Tiago Monteiro-Brás.

---

**Supplemental Figure 1.**
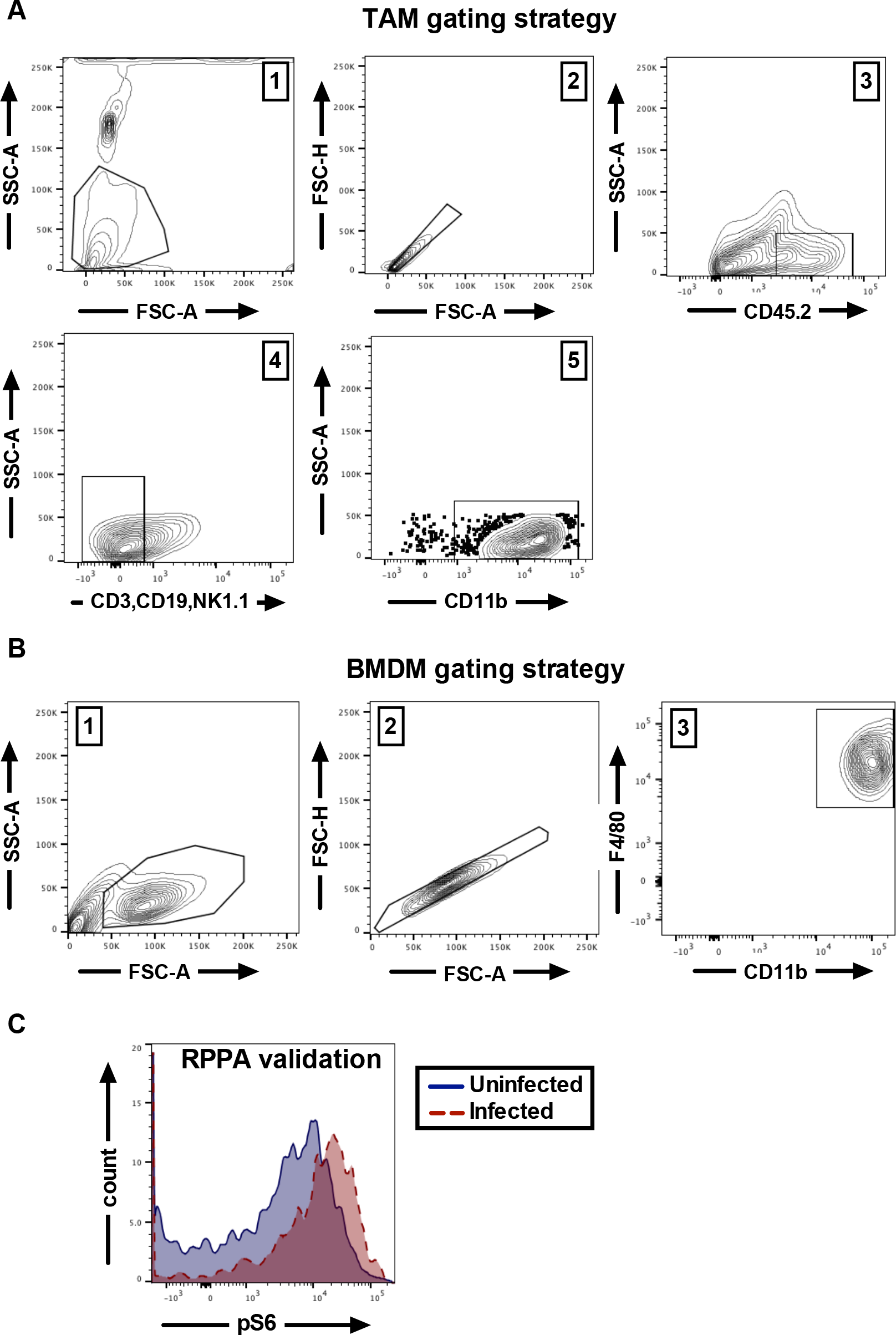
Representative gating strategies and RPPA validation **A)** Representative FACS plots from B16-F0 tumors indicating the gating strategy for assessing macrophage as shown in Figure 6C-D. Numbers indicate the order in which the gates were drawn. **B)** Representative FACS plots showing the gating strategy for assessment of BMDMs. Numbers indicate the order in which gates were drawn. **C)** Overlapping histograms of phospho-S6 (pS235-S236) in MCMV infected vs uninfected bone marrow macrophages, gated as in B.

**Supplemental Figure 2.**
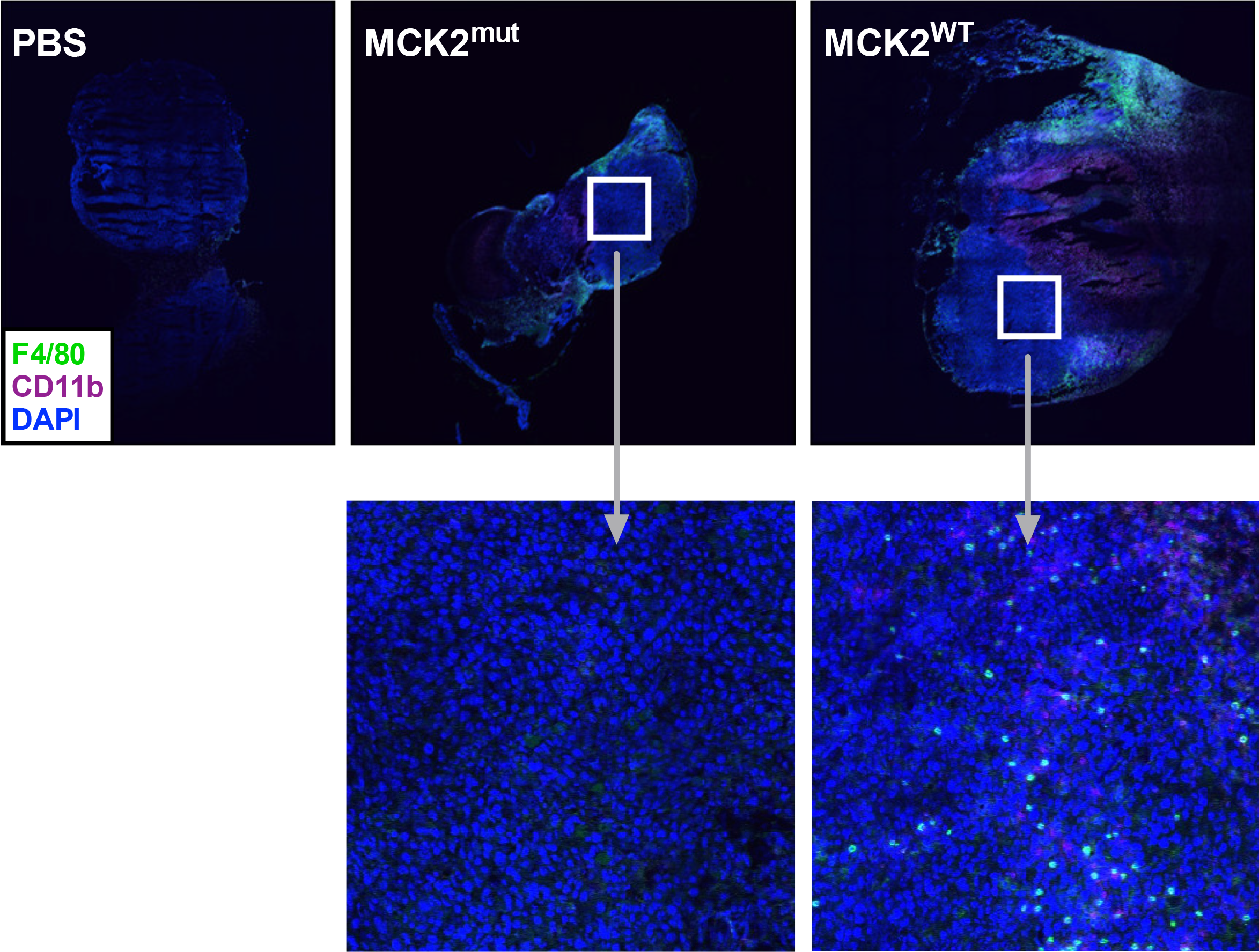
Macrophage infiltration into lesions is impaired without MCK2 CC function. Tiled images of a B16-F0 tumor treated IT with MCK2^mut^ virus, MCK2^WT^ virus, or PBS are shown. Tumors were excised on day 8 after the initial IT injection, frozen in O.C.T medium, and cryosectioned. All tumor sections were stained with F4/80, CD11b, and DAPI.

**Supplemental Table 1** Raw RPPA data.

## References

1. Crough, T. & Khanna, R. Immunobiology of Human Cytomegalovirus: from Bench to Bedside. Clin. Microbiol. Rev. 22, 76–98 (2009).

2. Manicklal, S., Emery, V. C., Lazzarotto, T., Boppana, S. B. & Gupta, R. K. The ‘silent’ global burden of congenital cytomegalovirus. Clin. Microbiol. Rev. 26, 86–102 (2013).

3. Karrer, U. et al. Memory inflation: continuous accumulation of antiviral CD8+ T cells over time. J. Immunol. Baltim. Md 1950 170, 2022–2029 (2003).

4. Ouyang, Q. et al. Age-associated accumulation of CMV-specific CD8+ T cells expressing the inhibitory killer cell lectin-like receptor G1 (KLRG1). Exp. Gerontol. 38, 911–920 (2003).

5. Sierro, S., Rothkopf, R. & Klenerman, P. Evolution of diverse antiviral CD8+ T cell populations after murine cytomegalovirus infection. Eur. J. Immunol. 35, 1113–1123 (2005).

6. Sylwester, A. W. et al. Broadly targeted human cytomegalovirus-specific CD4+ and CD8+ T cells dominate the memory compartments of exposed subjects. J. Exp. Med. 202, 673–685 (2005).

7. Snyder, C. M. et al. Memory Inflation During Chronic Viral Infection is Maintained by Continuous Production of Short-Lived Functional T Cells. Immunity 29, 650–659 (2008).

8. Smith, C. J., Turula, H. & Snyder, C. M. Systemic Hematogenous Maintenance of Memory Inflation by MCMV Infection. PLoS Pathog. 10, (2014).

9. Holtappels, R., Thomas, D., Podlech, J. & Reddehase, M. J. Two antigenic peptides from genes m123 and m164 of murine cytomegalovirus quantitatively dominate CD8 T-cell memory in the H-2d haplotype. J. Virol. 76, 151–164 (2002).

10. Snyder, C. M., Allan, J. E., Bonnett, E. L., Doom, C. M. & Hill, A. B. Cross-Presentation of a Spread-Defective MCMV Is Sufficient to Prime the Majority of Virus-Specific CD8+ T Cells. PLoS ONE 5, (2010).

11. Hansen, S. G. et al. Profound early control of highly pathogenic SIV by an effector memory T-cell vaccine. Nature 473, 523–527 (2011).

12. Hansen, S. G. et al. Cytomegalovirus vectors violate CD8+ T cell epitope recognition paradigms. Science 340, 1237874 (2013).

13. Hansen, S. G. et al. Immune clearance of highly pathogenic SIV infection. Nature 502, 100–104 (2013).

14. Hansen, S. G. et al. Effector memory T cell responses are associated with protection of rhesus monkeys from mucosal simian immunodeficiency virus challenge. Nat. Med. 15, 293–299 (2009).

15. Okoye, A. A. et al. Early antiretroviral therapy limits SIV reservoir establishment to delay or prevent post-treatment viral rebound. Nat. Med. 24, 1430–1440 (2018).

16. Xu, G., Smith, T., Grey, F. & Hill, A. B. Cytomegalovirus-based based cancer vaccines expressing TRP2 induce rejection of melanoma in mice. Biochem. Biophys. Res. Commun. 437, 287–291 (2013).

17. Qiu, Z. et al. Cytomegalovirus-Based Vaccine Expressing a Modified Tumor Antigen Induces Potent Tumor-Specific CD8(+) T-cell Response and Protects Mice from Melanoma. Cancer Immunol. Res. 3, 536–546 (2015).

18. Klyushnenkova, E. N. et al. A cytomegalovirus-based vaccine expressing a single tumor-specific CD8+ T cell epitope delays tumor growth in a murine model of prostate cancer. J. Immunother. Hagerstown Md 1997 35, 390–399 (2012).

19. Dekhtiarenko, I. et al. Peptide Processing Is Critical for T-Cell Memory Inflation and May Be Optimized to Improve Immune Protection by CMV-Based Vaccine Vectors. PLoS Pathog. 12, e1006072 (2016).

20. Beyranvand Nejad, E. et al. Demarcated thresholds of tumor-specific CD8 T cells elicited by MCMV-based vaccine vectors provide robust correlates of protection. J. Immunother. Cancer 7, (2019).

21. Grenier, J. M., Yeung, S. T., Qiu, Z., Jellison, E. R. & Khanna, K. M. Combining Adoptive Cell Therapy with Cytomegalovirus-Based Vaccine Is Protective against Solid Skin Tumors. Front. Immunol. 8, (2018).

22. Qiu, Z., Grenier, J. M. & Khanna, K. M. Reviving virus based cancer vaccines by using cytomegalovirus vectors expressing modified tumor antigens. Oncoimmunology 5, (2015).

23. Fukuhara, H., Ino, Y. & Todo, T. Oncolytic virus therapy: A new era of cancer treatment at dawn. Cancer Sci. 107, 1373–1379 (2016).

24. Eggermont, A. M. M., Crittenden, M. & Wargo, J. Combination Immunotherapy Development in Melanoma. Am. Soc. Clin. Oncol. Educ. Book 197–207 (2018). doi:10.1200/EDBK_201131

25. Chesney, J. et al. Randomized, Open-Label Phase II Study Evaluating the Efficacy and Safety of Talimogene Laherparepvec in Combination With Ipilimumab Versus Ipilimumab Alone in Patients With Advanced, Unresectable Melanoma. J. Clin. Oncol. 36, 1658–1667 (2018).

26. Tu, D.-G. et al. Salmonella inhibits tumor angiogenesis by downregulation of vascular endothelial growth factor. Oncotarget 7, 37513–37523 (2016).

27. Vola, M. et al. TLR7 agonist in combination with Salmonella as an effective antimelanoma immunotherapy. Immunotherapy 10, 665–679 (2018).

28. Quetglas, J. I. et al. Virotherapy with a Semliki Forest Virus-Based Vector Encoding IL12 Synergizes with PD-1/PD-L1 Blockade. Cancer Immunol. Res. 3, 449–454 (2015).

29. Sanchez-Paulete, A. R. et al. Intratumoral immunotherapy with XCL1 and sFlt3L encoded in recombinant Semliki Forest Virus-derived vectors fosters dendritic cell-mediated T cell cross-priming. Cancer Res. (2018). doi:10.1158/0008-5472.CAN-18-0933

30. Baird, J. R. et al. Immune-mediated regression of established B16F10 melanoma by intratumoral injection of attenuated Toxoplasma gondii protects against rechallenge. J. Immunol. Baltim. Md 1950 190, 469–478 (2013).

31. Fox, B. A., Sanders, K. L., Chen, S. & Bzik, D. J. Targeting tumors with nonreplicating Toxoplasma gondii uracil auxotroph vaccines. Trends Parasitol. 29, 431–437 (2013).

32. Dai, P. et al. Intratumoral delivery of inactivated modified vaccinia virus Ankara (iMVA) induces systemic antitumor immunity via STING and Batf3-dependent dendritic cells. Sci. Immunol. 2, (2017).

33. Michaelis, M., Doerr, H. W. & Cinatl, J. The Story of Human Cytomegalovirus and Cancer: Increasing Evidence and Open Questions. Neoplasia N. Y. N 11, 1–9 (2009).

34. Herbein, G. The Human Cytomegalovirus, from Oncomodulation to Oncogenesis. Viruses 10, (2018).

35. Forslund, O., Holmquist Mengelbier, L. & Gisselsson, D. Regarding human cytomegalovirus in neuroblastoma. Cancer Med. 3, 1038–1040 (2014).

36. Rådestad, A. F. et al. Impact of Human Cytomegalovirus Infection and its Immune Response on Survival of Patients with Ovarian Cancer. Transl. Oncol. 11, 1292–1300 (2018).

37. Zafiropoulos, A., Tsentelierou, E., Billiri, K. & Spandidos, D. A. Human herpes viruses in non-melanoma skin cancers. Cancer Lett. 198, 77–81 (2003).

38. Cobbs, C. S. et al. Human Cytomegalovirus Infection and Expression in Human Malignant Glioma. Cancer Res. 62, 3347–3350 (2002).

39. Harkins, L. et al. Specific localisation of human cytomegalovirus nucleic acids and proteins in human colorectal cancer. Lancet Lond. Engl. 360, 1557–1563 (2002).

40. Huang, E. S. & Roche, J. K. Cytomegalovirus D.N.A. and adenocarcinoma of the colon: Evidence for latent viral infection. Lancet Lond. Engl. 1, 957–960 (1978).

41. Hadaczek, P. et al. Cidofovir: A Novel Antitumor Agent for Glioblastoma. Clin. Cancer Res. Off. J. Am. Assoc. Cancer Res. 19, 6473–6483 (2013).

42. Stragliotto, G. et al. Effects of valganciclovir as an add-on therapy in patients with cytomegalovirus-positive glioblastoma: a randomized, double-blind, hypothesis-generating study. Int. J. Cancer 133, 1204–1213 (2013).

43. Crough, T. et al. Ex vivo functional analysis, expansion and adoptive transfer of cytomegalovirus-specific T-cells in patients with glioblastoma multiforme. Immunol. Cell Biol. 90, 872–880 (2012).

44. Bayer, C. et al. Human cytomegalovirus infection of M1 and M2 macrophages triggers inflammation and autologous T-cell proliferation. J. Virol. 87, 67–79 (2013).

45. Schuessler, A., Walker, D. G. & Khanna, R. Cytomegalovirus as a novel target for immunotherapy of glioblastoma multiforme. Front. Oncol. 4, 275 (2014).

46. Krenzlin, H. et al. Cytomegalovirus promotes murine glioblastoma growth via pericyte recruitment and angiogenesis. J. Clin. Invest. 130, (2019).

47. Price, R. L. et al. Cytomegalovirus Infection Leads to Pleomorphic Rhabdomyosarcomas in Trp53+/− Mice. Cancer Res. 72, 5669–5674 (2012).

48. Erlach, K. C., Böhm, V., Seckert, C. K., Reddehase, M. J. & Podlech, J. Lymphoma Cell Apoptosis in the Liver Induced by Distant Murine Cytomegalovirus Infection. J. Virol. 80, 4801–4819 (2006).

49. Erlach, K. C., Reddehase, M. J. & Podlech, J. Mechanism of tumor remission by cytomegalovirus in a murine lymphoma model: evidence for involvement of virally induced cellular interleukin-15. Med. Microbiol. Immunol. (Berl.) 204, 355–366 (2015).

50. Erlach, K. C., Podlech, J., Rojan, A. & Reddehase, M. J. Tumor Control in a Model of Bone Marrow Transplantation and Acute Liver-Infiltrating B-Cell Lymphoma: an Unpredicted Novel Function of Cytomegalovirus. J. Virol. 76, 2857–2870 (2002).

51. Erkes, D. A. et al. Intratumoral Infection with Murine Cytomegalovirus Synergizes with PD-L1 Blockade to Clear Melanoma Lesions and Induce Long-term Immunity. Mol. Ther. J. Am. Soc. Gene Ther. 24, 1444–1455 (2016).

52. Lepiller, Q., Abbas, W., Kumar, A., Tripathy, M. K. & Herbein, G. HCMV activates the IL-6-JAK-STAT3 axis in HepG2 cells and primary human hepatocytes. PloS One 8, e59591 (2013).

53. Kumar, A. et al. Tumor control by human cytomegalovirus in a murine model of hepatocellular carcinoma. Mol. Ther. Oncolytics 3, 16012 (2016).

54. Koffron, A. J. et al. Cellular Localization of Latent Murine Cytomegalovirus. J. Virol. 72, 95–103 (1998).

55. Hahn, G., Jores, R. & Mocarski, E. S. Cytomegalovirus remains latent in a common precursor of dendritic and myeloid cells. Proc. Natl. Acad. Sci. U. S. A. 95, 3937–3942 (1998).

56. Kondo, K., Xu, J. & Mocarski, E. S. Human cytomegalovirus latent gene expression in granulocyte-macrophage progenitors in culture and in seropositive individuals. Proc. Natl. Acad. Sci. U. S. A. 93, 11137–11142 (1996).

57. Kondo, K., Kaneshima, H. & Mocarski, E. S. Human cytomegalovirus latent infection of granulocyte-macrophage progenitors. Proc. Natl. Acad. Sci. U. S. A. 91, 11879–11883 (1994).

58. Pollock, J. L., Presti, R. M., Paetzold, S. & Virgin, H. W. Latent murine cytomegalovirus infection in macrophages. Virology 227, 168–179 (1997).

59. Taylor-Wiedeman, J., Sissons, J. G., Borysiewicz, L. K. & Sinclair, J. H. Monocytes are a major site of persistence of human cytomegalovirus in peripheral blood mononuclear cells. J. Gen. Virol. 72 (Pt 9), 2059–2064 (1991).

60. Stoddart, C. A. et al. Peripheral blood mononuclear phagocytes mediate dissemination of murine cytomegalovirus. J. Virol. 68, 6243–6253 (1994).

61. Gabrilovich, D. I., Ostrand-Rosenberg, S. & Bronte, V. Coordinated regulation of myeloid cells by tumours. Nat. Rev. Immunol. 12, 253–268 (2012).

62. Mantovani, A., Marchesi, F., Malesci, A., Laghi, L. & Allavena, P. Tumor-Associated Macrophages as Treatment Targets in Oncology. Nat. Rev. Clin. Oncol. 14, 399–416 (2017).

63. Chanmee, T., Ontong, P., Konno, K. & Itano, N. Tumor-Associated Macrophages as Major Players in the Tumor Microenvironment. Cancers 6, 1670–1690 (2014).

64. Jarosz-Biej, M. et al. M1-like macrophages change tumor blood vessels and microenvironment in murine melanoma. PloS One 13, e0191012 (2018).

65. Lyons, Y. A. et al. Macrophage depletion through colony stimulating factor 1 receptor pathway blockade overcomes adaptive resistance to anti-VEGF therapy. Oncotarget 8, 96496–96505 (2017).

66. Perry, C. J. et al. Myeloid-targeted immunotherapies act in synergy to induce inflammation and antitumor immunity. J. Exp. Med. 215, 877–893 (2018).

67. Hoves, S. et al. Rapid activation of tumor-associated macrophages boosts preexisting tumor immunity. J. Exp. Med. 215, 859–876 (2018).

68. Overwijk, W. W. & Restifo, N. P. B16 as a Mouse Model for Human Melanoma. Curr. Protoc. Immunol. Ed. John E Coligan Al CHAPTER, Unit-20.1 (2001).

69. Noda, S. et al. Cytomegalovirus MCK-2 controls mobilization and recruitment of myeloid progenitor cells to facilitate dissemination. Blood 107, 30–38 (2006).

70. Zurbach, K. A., Moghbeli, T. & Snyder, C. M. Resolving the titer of murine cytomegalovirus by plaque assay using the M2-10B4 cell line and a low viscosity overlay. Virol. J. 11, 71 (2014).

71. Reddehase, M. J., Fibi, M. R., Keil, G. M. & Koszinowski, U. H. Late-phase expression of a murine cytomegalovirus immediate-early antigen recognized by cytolytic T lymphocytes. J. Virol. 60, 1125–1129 (1986).

72. Shi, W., Oshlack, A. & Smyth, G. K. Optimizing the noise versus bias trade-off for Illumina whole genome expression BeadChips. Nucleic Acids Res. 38, e204–e204 (2010).

73. Trapnell, C. et al. The dynamics and regulators of cell fate decisions are revealed by pseudotemporal ordering of single cells. Nat. Biotechnol. 32, 381–386 (2014).

74. Qiu, X. et al. Single-cell mRNA quantification and differential analysis with Census. Nat. Methods 14, 309–315 (2017).

75. Qiu, X. et al. Reversed graph embedding resolves complex single-cell trajectories. Nat. Methods 14, 979–982 (2017).

76. Erkes, D. A. et al. Intratumoral Infection with Murine Cytomegalovirus Synergizes with PD-L1 Blockade to Clear Melanoma Lesions and Induce Long-term Immunity. Mol. Ther. J. Am. Soc. Gene Ther. 24, 1444–1455 (2016).

77. Daley-Bauer, L. P., Roback, L. J., Wynn, G. M. & Mocarski, E. S. Cytomegalovirus hijacks CX3CR1(hi) patrolling monocytes as immune-privileged vehicles for dissemination in mice. Cell Host Microbe 15, 351–362 (2014).

78. Saederup, N., Aguirre, S. A., Sparer, T. E., Bouley, D. M. & Mocarski, E. S. Murine Cytomegalovirus CC Chemokine Homolog MCK-2 (m131-129) Is a Determinant of Dissemination That Increases Inflammation at Initial Sites of Infection. J. Virol. 75, 9966–9976 (2001).

79. Wagner, F. M. et al. The viral chemokine MCK-2 of murine cytomegalovirus promotes infection as part of a gH/gL/MCK-2 complex. PLoS Pathog. 9, e1003493 (2013).

80. Kudchodkar, S. B., Yu, Y., Maguire, T. G. & Alwine, J. C. Human Cytomegalovirus Infection Induces Rapamycin-Insensitive Phosphorylation of Downstream Effectors of mTOR Kinase. J. Virol. 78, 11030–11039 (2004).

81. Vomaske, J. et al. Cytomegalovirus CC Chemokine Promotes Immune Cell Migration. J. Virol. 86, 11833–11844 (2012).

82. Chan, G., Bivins-Smith, E. R., Smith, M. S., Smith, P. M. & Yurochko, A. D. Transcriptome analysis reveals human cytomegalovirus reprograms monocyte differentiation toward an M1 macrophage. J. Immunol. Baltim. Md 1950 181, 698–711 (2008).

83. Metcalfe, W., Anderson, J., Trinh, V. A. & Hwu, W.-J. Anti-programmed cell death-1 (PD-1) monoclonal antibodies in treating advanced melanoma. Discov. Med. 19, 393–401 (2015).

84. Forbes, N. S. et al. White paper on microbial anti-cancer therapy and prevention. J. Immunother. Cancer 6, 78 (2018).

85. Ma, H. S. et al. A CD40 Agonist and PD-1 Antagonist Antibody Reprogram the Microenvironment of Nonimmunogenic Tumors to Allow T-cell–Mediated Anticancer Activity. Cancer Immunol. Res. 7, 428–442 (2019).

